# Network Structure During Encoding Predicts Working Memory Performance

**DOI:** 10.1101/409615

**Authors:** Anirudh Wodeyar, Ramesh Srinivasan

## Abstract

Working memory operates through networks that integrate distributed modular brain activity. We characterize the structure of networks in different electroencephalographic frequency bands while individuals perform a working memory task. The objective was to identify network properties that support working memory function during the encoding, maintenance, and retrieval of memory. In each EEG frequency band, we estimated a complex-valued Gaussian graphical model to characterize the structure of brain networks using measures from graph theory. Critically, the structural characteristics of brain networks that facilitate performance are all established during encoding, suggesting that they reflect the effect of attention on the quality of the representation in working memory. Segregation of networks in the alpha and beta bands during encoding increased with accuracy. In the theta band, greater integration of functional clusters involving the temporal lobe with other cortical areas predicted faster response time, starting in the encoding interval and persisting throughout the task, indicating that functional clustering facilitates rapid memory manipulation.

## Introduction

The brain’s persistent representation of items retained in working memory (an active short term store) has been investigated through neuroimaging and electrophysiology^1–4^. Maintenance of memories has long been believed to be managed by elevated activity (increased firing rate) in the pre-frontal cortex^5^. Pre-frontal cortex orients attention towards representations maintained in the brain. Subsequent studies have found evidence for representation of information during maintenance in sensory, parietal, temporal and frontal areas^1–4, 6^ with frontal areas retaining information at a more abstract scale than sensory areas^7^. This distributed representation of information can be expected for a self organized system^8, 9^ such as the brain that integrates specialized modules into a functional network.

The structure of the distributed system during behavior can be examined through measuring network topology using the analytic framework of graph theory. Complex networks observed in brain structure and function^10–14^ display short network distances between areas (short path length) and local networks that are strongly interconnected (high clustering coefficient). This indicates the brain is optimally functionally integrated while also maintaining functional segregation through minimal connections between specialized modules permitting efficient information transfer. A number of studies have shown that modulation of the brain networks’ functional integration and segregation plays a role in cognition and disease^15, 16^.

Working memory capacity and performance have been associated with properties of global network topology in functional networks estimated through electroencephalography (EEG), magnetoencephalography(MEG), functional magnetic resonance imaging (fMRI) and in structural networks estimated using diffusion magnetic resonance imaging (dMRI). In an MEG study, an estimate of the overall functional segregation (modularity) in the beta frequency band was negatively correlated to levels of an N-Back task^17^, i.e., increasing levels of the N-back task lead to increased global functional integration. Working memory load was also associated with increased clustering coefficient (increased functional clustering) in alpha/beta bands in another MEG study^18^. In a fMRI study, greater global functional integration of networks was associated with greater working memory capacity^19^. Further, intra-individual variation in capacity across two different sessions was explained by increases in global functional integration. Working memory training has been shown to cause large scale network reconfiguration in networks estimated using source-localized EEG and structural networks estimated with dMRI^20, 21^. Greater efficiency of information transfer as revealed by a higher small-worldness coefficient (an estimate of the efficiency of information transfer in the network) of EEG source localized theta band networks was correlated with improved capacity^20^. Structural connectivity estimates showed that greater efficiency of information transfer in the fronto-parietal attention network was associated with improvement in capacity during working memory training^21^.

While working memory capacity is clearly impacted by network structure, there is limited research on the influence of dynamic characteristics of these networks during working memory processing on performance (accuracy/reaction time). It is assumed that network activity is related to maintenance of memory but this has not been specifically related to performance. In our study, we use a task where the trial structure separates the operations of encoding, maintenance and retrieval into different intervals. Attention to the stimulus is critical to the encoding period^22^ where brain networks are established that are engaged in retaining a memory during the maintenance interval. This task paradigm with separate encoding, maintenance and retrieval has been shown to engage networks in different frequency bands, alpha and beta for attention and theta and gamma for memory^23–27^.

We used a graphical model of effective connectivity (complex gaussian graphical model) in each EEG frequency band to investigate the relationship between network structure and performance in a working memory task. We characterized the network structure in each frequency band using: (1) connection density (mean degrees), (2) integration of functional clusters in the network (clustering coefficient) and (3) a global measure of functional segregation (path length). We hypothesized that network characteristics as estimated through graph theoretic measures would be correlated with accuracy and response time revealing the important functional properties of network topology.

## Methods

### Participants

Seventeen healthy volunteers participated in the study (12 male, 5 female). All subjects were right handed and had normal or corrected vision. Participants did not report any neurological disorders. Each participant was compensated $30 for 1.5 hours. The study was approved by the University of California, Irvine institutional review board. The experiment was conducted according to the approved guidelines. All participants provided written informed consent.

### Working Memory Task

Working memory experiments typically follow an encoding, delay and retrieval paradigm. In our experiment participants had to remember the orientations of two Gabor patches embedded in noise over a delay period of random length before being tested on their recollection of the orientation of one of the Gabors (shown in Figure 1). Two Gabors of different orientations were presented, one on each side of the screen. The Gabor on the left and right were flickered at 24 and 40 Hz respectively, while background noise changed at 30 Hz. The response to the flickers is not used in the analysis covered in this paper. After an encoding period of 1 second, participants retained the orientations of both Gabors for a random delay (between 1 to 3.5 seconds) while changing noise continued on the screen. After the delay, a probe Gabor was displayed either on the right or left side of the screen and rotated 30 degrees from the original orientation while flickering at the same rate as the initially presented Gabor. Participants needed to respond as to whether the Gabor presented as a probe had been rotated clockwise or counter-clockwise within a 2 second interval after the probe was presented. Participants performed 400 trials of the task.

**Figure 1.**
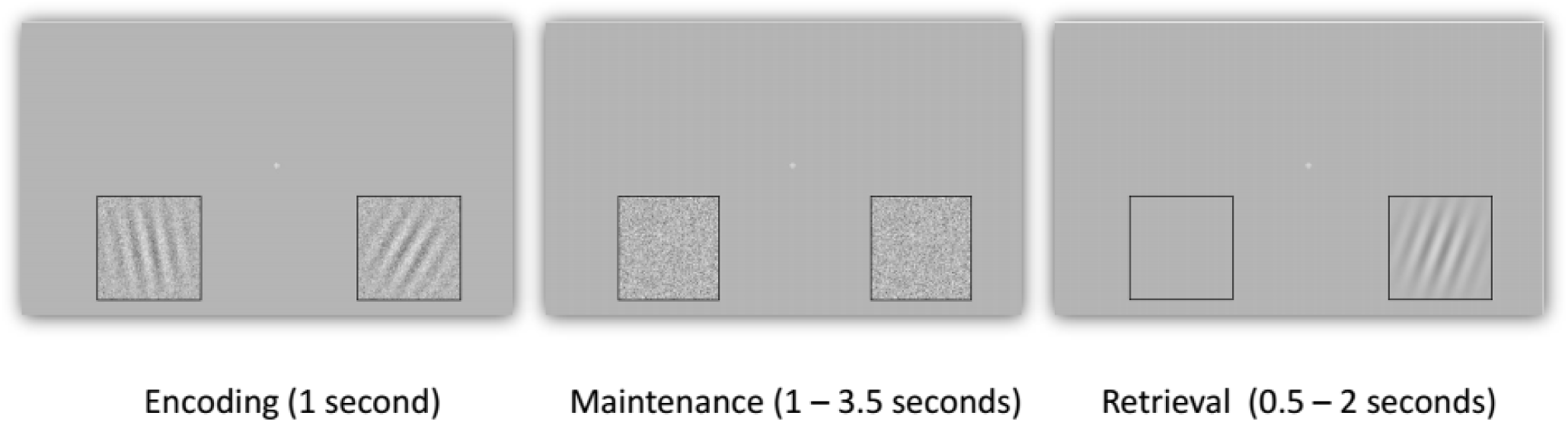
Experimental stimulus: Participants in the task fixated on the white cross. Each trial began after a uniformly random interval of time, and each delay (maintenance) was also uniformly distributed between 1 to 3.5 seconds. Participants needed to remember the orientation of the two Gabors on either side of the screen for a random delay before a probe appeared. The probe was rotated 30 degrees relative to the original stimulus clockwise or counter-clockwise, and participants needed to respond which way it was rotated. We took the 500 ms prior to the start of the trial, the encoding period, the first second of the maintenance period, and the first 500 ms of the retrieval interval and analyzed network properties during this period of time.

### EEG Recording and Preprocessing

We collected EEG from 128 channels using the Electronic Geodesics (EGI) hydrocel sensor net (HSN). The EEG was sampled at a rate of 1000 Hz, and hardware high pass and low pass filtered at 0.1 Hz and 50 Hz. EEG data was first detrended (Matlab, Natick, MA). Data was then cleaned of gross movement artifacts manually and common average referenced. An independent components analysis (ICA) was performed on each participant’s data to remove eye blink and muscle artifact components. We used the cleaned data from 15 participants who yielded 300 trials. Two participants were removed since they had fewer than 300 artifact-free trials.

### Sliding Window Fourier Analysis

We performed graph theoretical analysis of the dynamic structures present over the pre-encoding (500ms), encoding (1 sec), maintenance (1 sec) and retrieval (500 ms) periods. The frequency bands were defined as: theta (5-7 Hz), alpha (8-12 Hz), beta (13-29 Hz excluding flicker frequencies of 24 Hz) and gamma (31 - 50 Hz excluding flicker frequency of 40 Hz). Preprocessed EEG time series data was transformed to the frequency domain using a fast fourier transform (Matlab) in sliding 500 millisecond segments that were moved forward 100 ms at a time.

### Complex Gaussian Graphical Models

We frame the problem of estimating effective connectivity in terms of the complex valued data after Fourier transform of every trial from each electrode. Every epoch (any one trial) of the EEG data gives us a realization of the complex multivariate gaussian random process in any one frequency band.

We define the complex vectors *z* = *x* + *iy* and *z*^*H*^ = *x* − *iy*, and the complex augmented vector *Z* for a specific frequency *Ω*, for any one epoch, at all electrodes *C*. Using *Z* = [*z, z*^*H*^] and 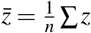, we define the complex multivariate normal (CMVN) for a complex Gaussian process^28^ over the electrodes as:

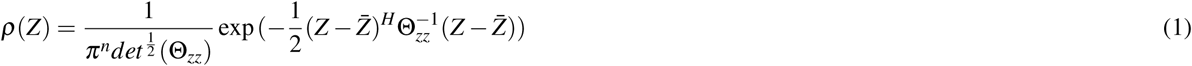

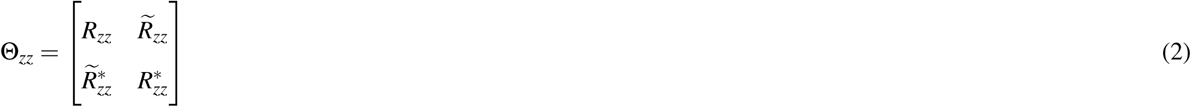

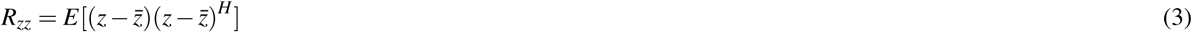

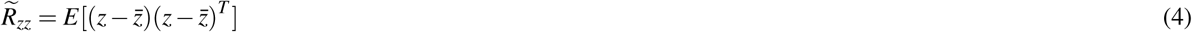

We subtract 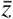 to remove any non-stationary mean values (i.e., evoked potentials) before computing covariance and complementary covariance. Any value in the precision matrix is an estimate of the conditional covariance 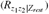 between any two variables (here electrodes) given the other variables^29^. When the value for precision is zero in the Schur complements of both the covariance and complementary covariance matrix, we can state that those electrodes are conditionally independent under the CMVN. By establishing the framework of the CMVN, we possess the ability to estimate which pairs of electrodes are in fact conditionally independent, and by its complement, which are conditionally dependent. The set of conditional dependencies among variables defines a complex-Gaussian graphical model of the EEG data, where the edges reflect effective connectivity between electrodes. Estimating conditional dependence reduces the problem of common inputs (and by extension false positive connections)^30^. This allows for a better approximation to the definition of an edge (connection) needed for graph theoretical analysis^31^.

### Graphical Lasso

The graphical lasso^32^ optimizes the multivariate normal likelihood function with added L1 norm penalization to estimate the precision matrix. It assumes a sparse connectivity structure or that there are zero valued entries present in majority in the precision matrix. In this way, we allow our estimates to be more robust for the precision values that are retained. In order to apply the lasso, we optimize over the following function to estimate the precision:

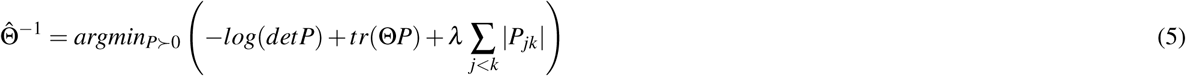

The parameter *λ*determines the cachet of precision values retained through the process of optimization. Using simulations of networks with different number of edges, we found that a *λ*= 0.1 * *max*(*Covariance*) is appropriate to maximize accuracy and minimize common input effects. We also looked at penalizations of 0.05 and 0.2 and found qualitatively similar results.

### Volume Conduction Model

In order to show how using the cGGM suppresses effects of volume conduction we made use of a simulation of volume conduction in a realistic head model. We generated an estimate of the boundary of these layers from an magnetic resonance image (MRI) of the fsaverage brain^33^. We made use of these surfaces to build a boundary element model of the brain, skull and scalp^34^. Conductivities of the tissue compartments were obtained from the literature^35^. In order to make use of a realistic distribution of sources, a surface was fit to each hemisphere of the cortex using Freesurfer. We reduce the 81k vertices of the cortex to 142 sources based on a modified Desikan-Killiany atlas^36^. Each source was represented by the central vertex of the surface patch of the source.

We simulated the effects of volume conduction by assuming each source is an independent complex-Gaussian random process. Within our simulated cortical surface all precision values are zero and there are no connections. At the electrodes, the ideal estimator of connectivity should be an identity matrix, showing the complete disconnection within the brain. Given the lead field matrix *L* of dimension *N*_*e*_*xN*_*s*_ where *N*_*e*_ is number of electrodes and *N*_*s*_ is number of sources, the relationship between the source cross-spectrum *R*_*s*_ and the electrode cross spectrum *R*_*e*_ is given as:

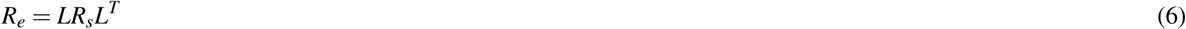

In our simulation *R*_*s*_ = *I* or in other words, we have an independent normal source at each point on the brain mesh and any non-zero values in the cross-spectrum calculated from the signals at the electrodes reflects only volume conduction. We compared our cGGM estimate of the edges of the graphical model to a simple threshold of the coherence values as is commonly used to define connections for graph theoretic analyses^37, 38^. Figure 2 shows the false positives in the cGGM over a range of penalization compared to using the same value as a threshold for the coherence. At all choices the cGGM results in far fewer false edges. Some volume conduction is always present in the EEG data and simple threshold on coherence is less effective at removing volume conduction^39^ than the cGGM model of effective connectivity.

**Figure 2.**
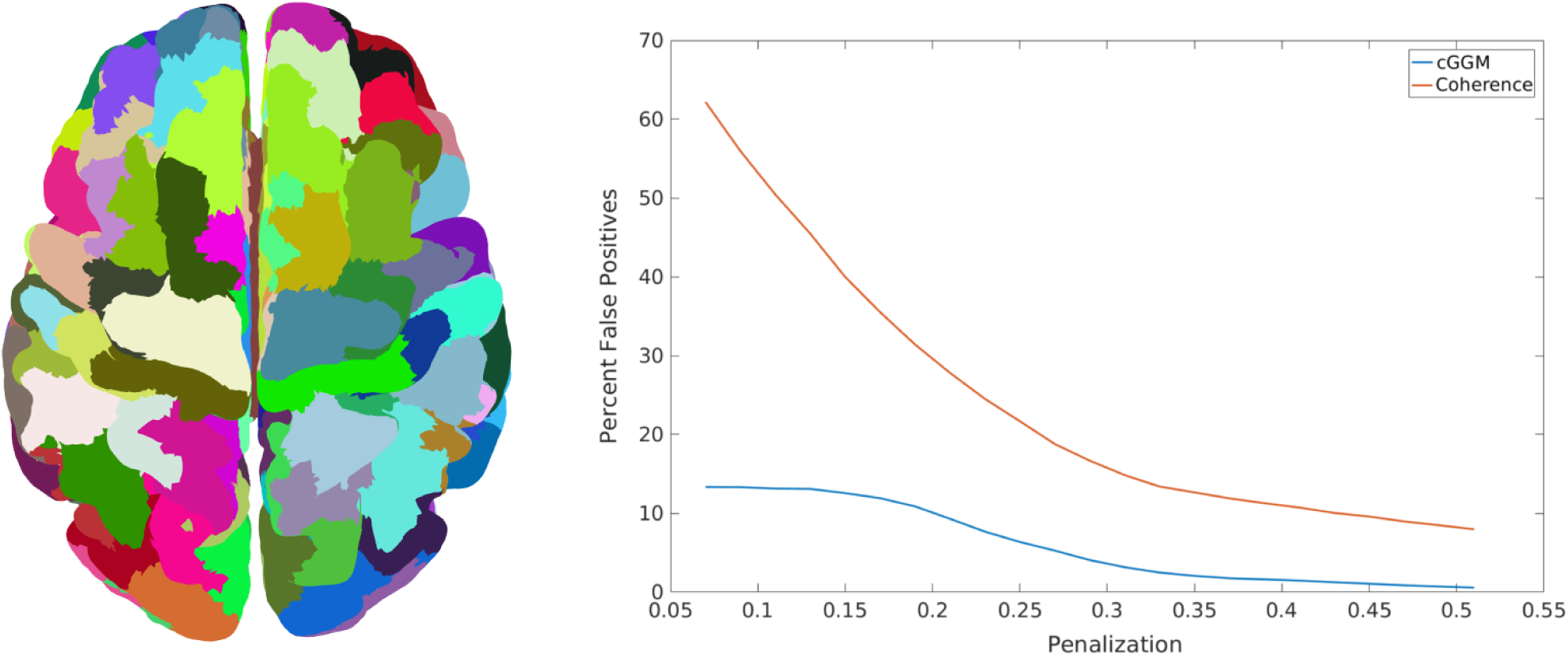
Volume Conduction Simulation: We used 142 sources from a modified Desikan-Killiany atlas (depicted on the left) and simulated activity at the scalp assuming each is a standard normal independent source. Using this simulated data, we estimated the number of connections present between electrodes when estimating coherence and the cGGM model. Percent of false positive connections (relative to total connections possible) were calculated across a range of penalization values for the graphical lasso for the cGGM model. We thresholded the coherence at the same penalization value. We can see that the cGGM model is very effective at reducing the number of false positives due to volume conduction while thresholding the coherence is inadequate for removing volume conduction effects.

### Graph Theory

We define the networks estimated by the complex-Gaussian graphical model by using all non-zero edges discovered from the graphical lasso. We estimate graph theoretic metrics using the Brain Connectivity Toolbox^40^. We make use of three measures in this study:

1. The basic measure of a network is the **degree** of a node (connectivity), that is, the number of edges a node shares with other nodes in an undirected graph. This measures how many nodes can communicate with any given node.
2. The local integration of functional clusters of the network is measured by the **clustering coefficient** (CC). The CC is the ratio of the number of triangles to number of triples in a network and helps give an estimate of how often the nodes one degree apart are connected to one another. 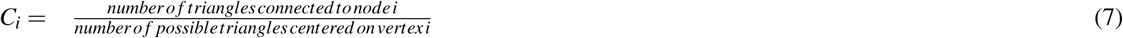

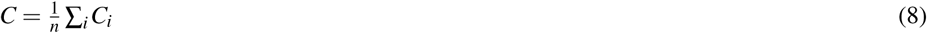
3. A metric that describes the global segregation of the network is the **path length**. Path length (L) is the average shortest path (*d*) between all pairs of nodes. 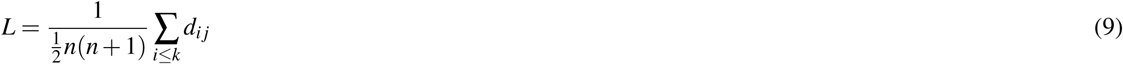

A higher path length (larger L) indicates greater functional segregation. over the whole network.

These graph-theoretic metrics are inflated (degrees and clustering coefficient), or cannot be estimated correctly (path length) if common inputs influence the estimation of the original network^31^, such as with a network estimated using estimates of coherence between EEG channels. By using the cGGM we reduce the impact of common inputs and volume conduction by estimating the conditional dependence between electrodes. We can then use these graph theoretic metrics to look at the impact of connection density, local integration of functional clusters and the global segregation of networks in different EEG frequency bands on the working memory performance.

### Statistics

All statistics were estimated using functions in Matlab (Natick, MA). Chi-squared tests were used to determine the p-values of correlation between graph theoretic metrics and behavior before they were subjected to the false discovery rate or FDR^41^ to correct for both multiple time point testing and for testing multiple frequency bands. Wilcoxon rank-sum tests were used to test for differences between high and low accuracy groups and also between fast and slow responders. Chi-squared tests were also used to identify significantly higher presence of an edge in fast responders relative to slow responders. For all cases where correcting for multiple comparisons, we used the FDR and limited false positives to 5 percent of tests.

## Results

### Behavioral Results

Participants responded whether the probe Gabor was rotated clockwise or counter-clockwise within 2 seconds with reasonable ease. The mean reaction time for participants was 0.987 ± 0.187, and mean accuracy was 0.734 ± 0.086. The low average accuracy is possibly due to the distance between probe and stimulus being 30 degrees which is difficult to discriminate. In order to characterize the relationship between brain network topology and behavior, we created a median split between slow and fast responders and separately between low and high accuracy participants. The slow responders (high reaction time) were defined as having greater than median reaction time (1.1525 seconds ± 0.13) and fast responders (low reaction time) as having less than median reaction time (0.8537 seconds ± 0.1). The slow and fast responders had significantly different reaction times (Wilcoxon rank-sum test, p<0.001). Low accuracy participants had accuracy of 0.676 ± 0.04 while high accuracy participants had an accuracy around 0.8 ± 0.08. Low accuracy and high accuracy groups were significantly different (Wilcoxon rank-sum test, p<0.001).

### Functional Clustering in Theta Band and Reaction Time

Clustering coefficient is estimated as the ratio between number of triangular edge structures (three nodes A, B and C are connected to one another) and total possible triangles, and thus reflects how much functional clustering can be found in a sparse network. Clustering coefficient in the theta band has a consistent negative correlation to reaction time (p<0.05 corrected) over the entire duration of the trial as seen in Figure 3a. Greater functional clustering was correlated to faster response times. We performed a median split of the data and examined clustering coefficient averaged over the encoding interval in different areas of the brain. (Figure 3b). Clustering coefficient is reduced in the slow responders relative to fast responders during encoding in the temporal lobe (t(13) = 2.9886, p=0.0105). During the interval where correlation to behavior was highest (600-1100ms post stimulus onset, see Figure 5 for networks involved) we explored which connections influenced the higher average clustering coefficient in the temporal lobe. Clustering coefficient increases when a triangle is formed between three nodes of a network so we examined all occurrences of these triangles in fast responders relative to slow responders when at least one of the three nodes was an electrode over the temporal lobe. We found that there are frontal to parietal/occipital connections (where one temporal node connects to both a frontal and a parietal/occipital node) and temporo-occipital connections (where two temporal nodes connect to an occipital node), as shown in Figure 3c. These connections are responsible for the increased clustering coefficient (Chi-squared test of proportions, p< 0.05 uncorrected) among faster responders. Functional clustering between temporal lobe and the rest of the brain is likely involved in sustaining representations in the brain in the focus of attention. Faster responders, by having more functional clustering have more functional integration between temporal lobe and the occipito-parietal areas.

**Figure 3.**
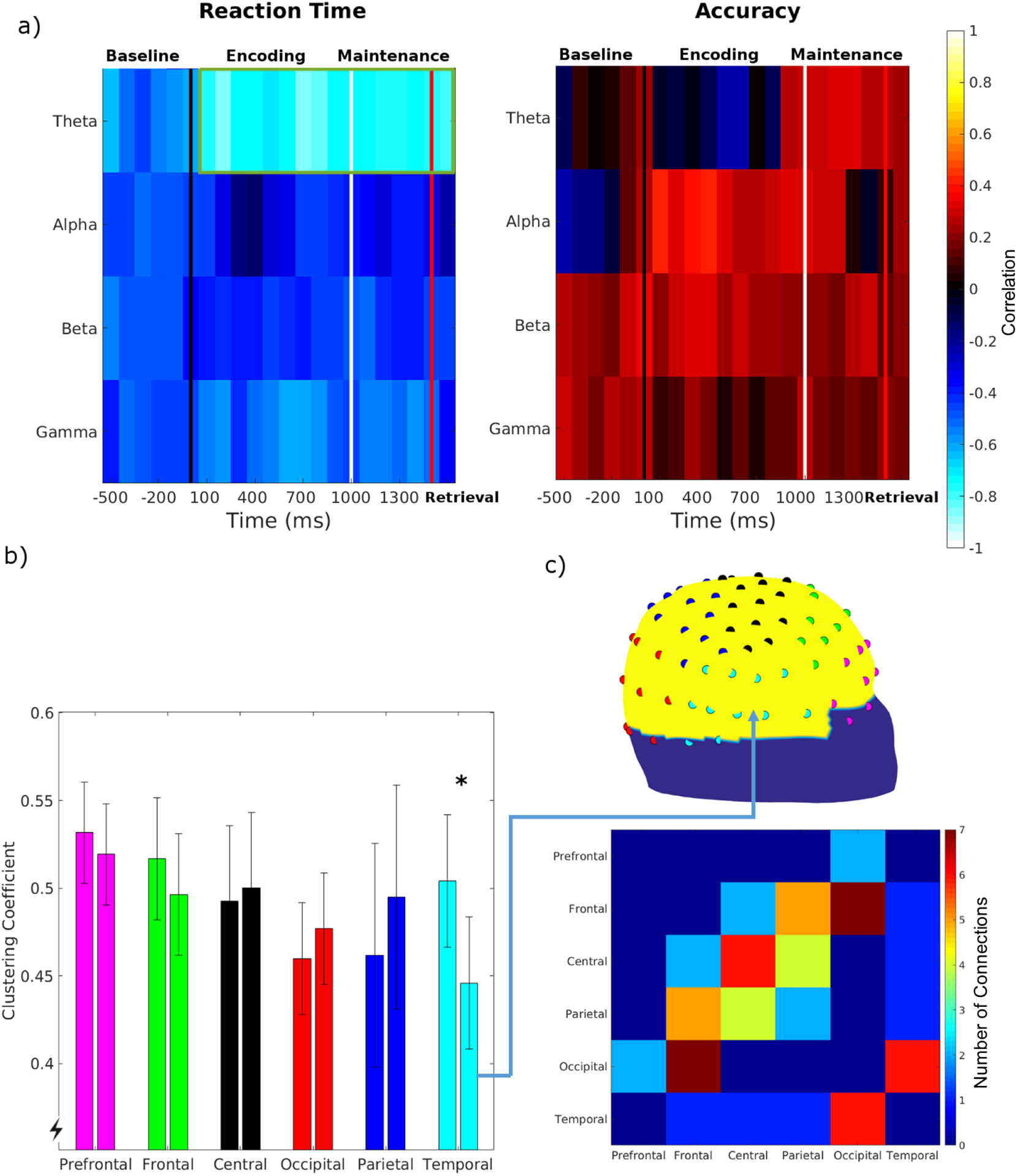
Functional Clustering and Performance: (a) Correlation between clustering coefficient in different frequency bands and reaction time (left) and accuracy (right). Theta band clustering coefficient is correlated (p<0.05, FDR corrected) to reaction time through the entire trial period. In the lower right figure (b) we show the difference in clustering coefficients between fast (left) and slow (right) responders, there is a significantly higher local integration of functional clusters formed by the temporal lobe (p=0.0105) with the rest of the brain for the fast responders. The figure on the right (in (c) upper image) shows which electrodes were chosen for represent activity from different areas and the lower image shows the number of connections per electrode between different areas that supported functional clustering with the temporal lobe. These are the connections that are present significantly (p<0.05) more often in fast responders relative to slow responders and form one edge of the triangles formed by the temporal lobe.

**Figure 4.**
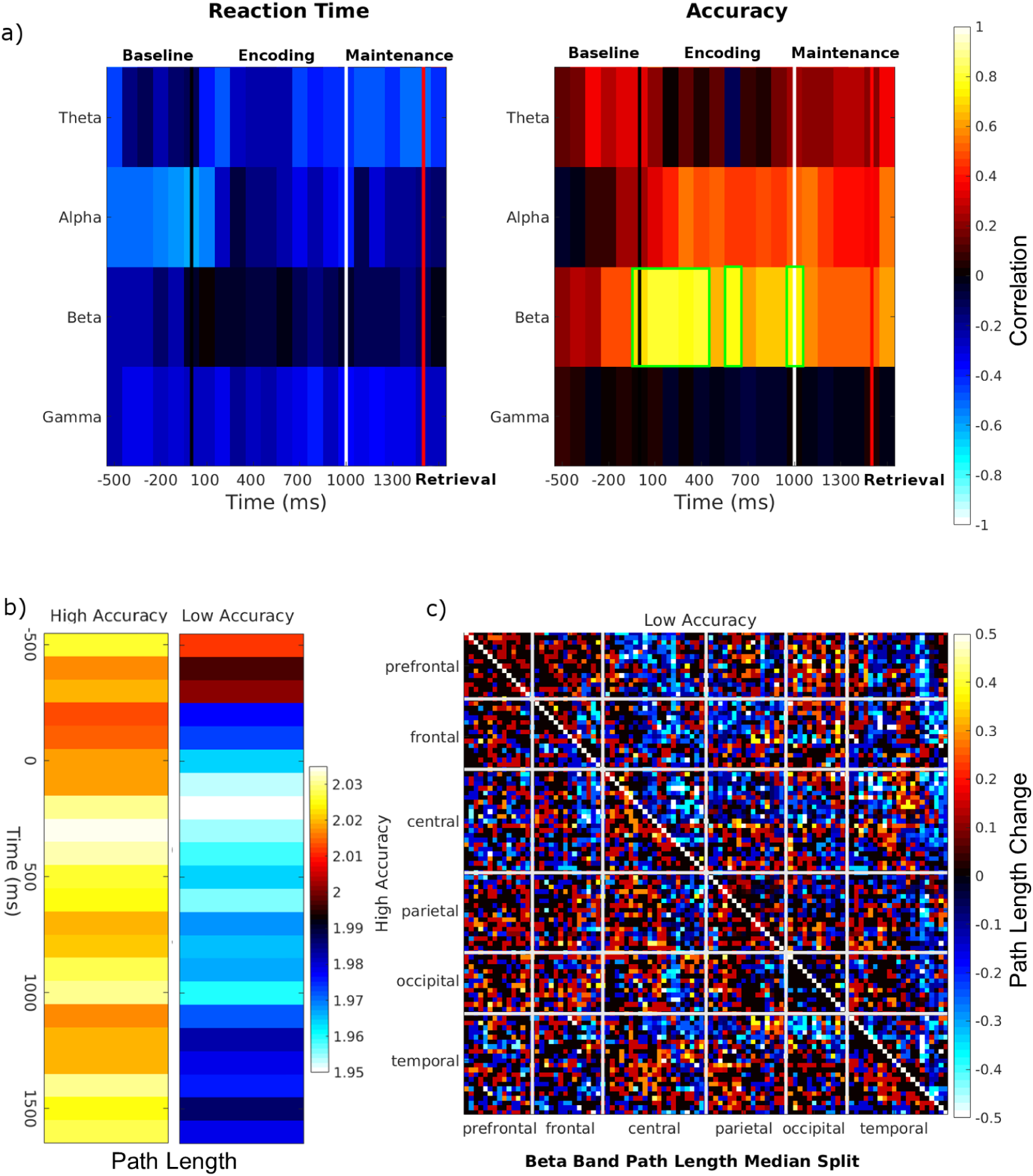
Functional Segregation and Performance: (a) Correlation between the reaction time and the path length in different frequency bands on the left and the correlation between path length and accuracy on the right. Significant correlations between beta band path length and accuracy (p<0.05, FDR corrected) exist during encoding interval and are indicated by the green rectangles. Subjects were split based on median accuracy into high and low accuracy groups. In (b) is the averaged dynamics of path length change over time in the high and low accuracy groups. (c) Difference in path length between pairs of electrodes between 300 ms post stimulus presentation and the baseline averaged over the high and low accuracy groups. The upper triangle (above the diagonal) in (c) is the averaged difference in path length for low accuracy subjects while the lower triangle represents the high accuracy. There is a reduction in path length between temporal and central to other brain areas for low accuracy but not for high accuracy. In high accuracy subjects there is a general increase in path length between most brain areas.

**Figure 5.**
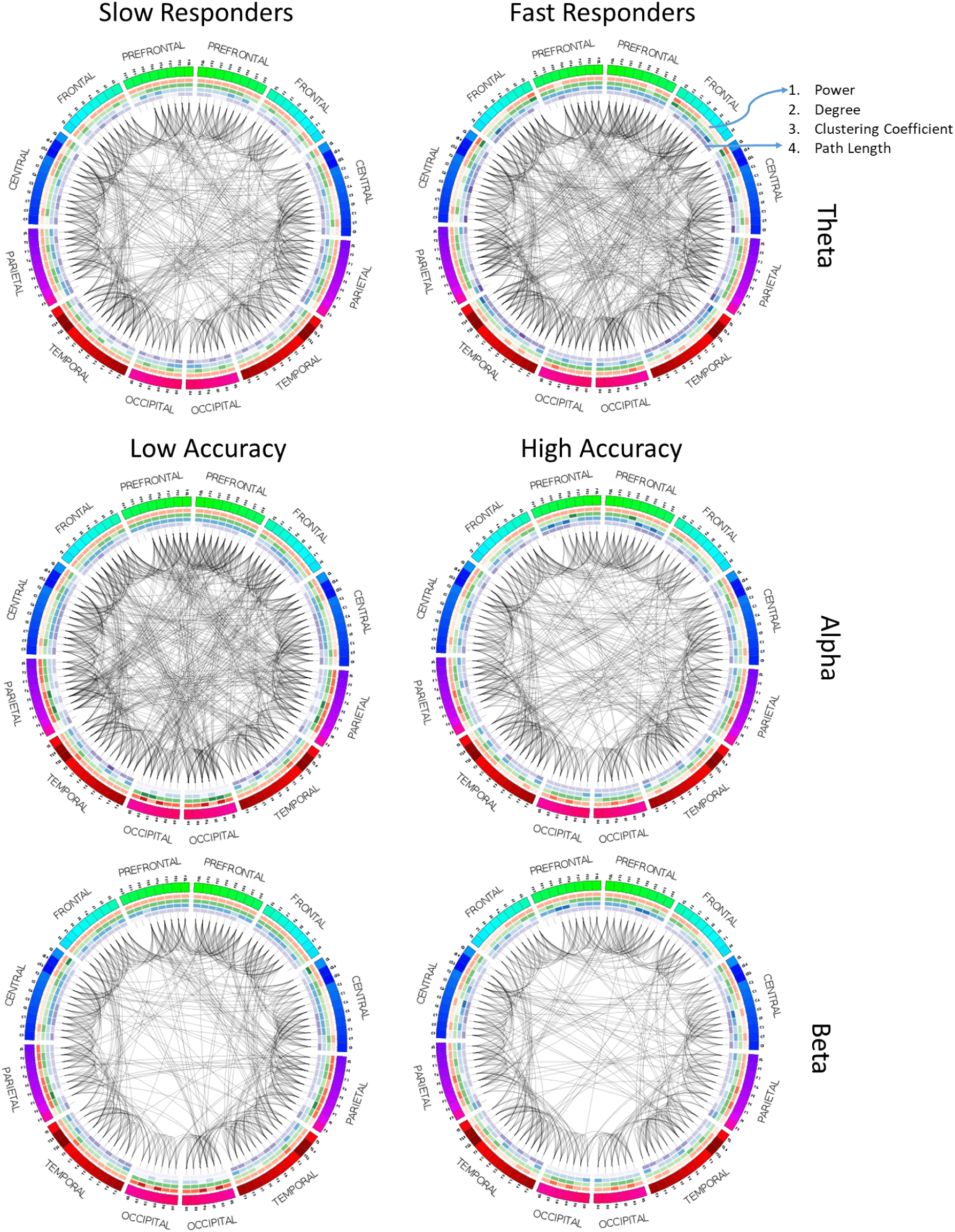
Graphical Models of Networks: We represent the networks formed in at least 4 participants or more among fast and slow responders in the theta band during the encoding interval (600-1100 ms). In the second row are networks among low accuracy and high accuracy participants in the alpha band during encoding (100 - 600ms). The bottom row depicts networks among low and high accuracy participants in the beta band during the encoding (200-700ms). We can see increased integration in theta and alpha bands networks while in beta band networks there is increased segregation in the network. In the heatmaps in outer rings of each circle plot, we represent the averaged power, degree, clustering coefficient and path length over all slow/fast responders or low/high accuracy participants.

### Beta Band Path Length and Accuracy

Path length is an estimate of the number of connections needed to get from one node to another. The path length of a network is computed by determining the shortest path length between each pair of nodes and averaging over all possible pairs of nodes. We tested each frequency band and time point and found that beta band path length is strongly positively correlated (p<0.05 FDR corrected) to accuracy during encoding (Figure 4a). Higher network path length indicates less direct connectivity and greater functional segregation. During encoding, greater functional segregation between brain areas in the beta band is related to higher accuracy. By reducing the number of direct connections between brain areas, there is reduced synchronization of activity during the encoding period permitting a greater diversity of activity (functional segregation). Functional segregation potentially creates more robust representations of the stimulus by allowing for redundancy without interference^7^.

We analyzed the data of low and high accuracy participants (based on median split of accuracy) separately. In Figure 4b, we see that the dynamics of path length change are different between high accuracy and low accuracy groups, with significant differences in path length (p < 0.05) during the encoding interval (−200 ms to 500 ms and 400 - 900 ms) between the two groups. The low accuracy group reaches the minimum path length at 200 ms into the encoding interval while the high accuracy group reaches maximum path length 300 ms into the encoding interval. We calculated the difference in path length for every pair of electrodes between the encoding period (at 300 ms during period of maximum correlation to behavior) and baseline and averaged it across high and low accuracy subjects (Figure 4c). Low accuracy subjects have reduced path length during encoding, particularly in temporal connections to the rest of the brain, and central connections to the occipital, parietal and temporal areas. High accuracy subjects show a broad increase in path length indicating global segregation. In Figure 5 (bottom row) we show the beta band networks in low and high accuracy participants.

### Alpha Connection Density and Accuracy

We calculated the mean degrees as an overall measure of connection density for each participant over the baseline, encoding, maintenance and retrieval periods by taking the average over all electrodes’ degrees calculated separately for each frequency band. We found that the connectivity was uncorrelated with reaction time in all frequency bands and positively correlated with accuracy only in the alpha band. Higher connection density (Figure 5) implied reduced accuracy (p<0.05, however this was not significant when FDR corrected) during the encoding period from 100 ms to 700 ms post stimulus onset, first 200 ms during maintenance and during the first 500 ms of retrieval. Alpha band desynchronization is a commonly reported phenomena upon the presentation of a visual stimulus^42^ and reflects the capture of the focus of attention by a stimulus^43^. Examining the individual subjects, we find that a subset of the subjects actively increased connection density (higher average degrees) in the alpha band during the encoding interval and exhibited lower accuracy. Participants who enhance connectivity (have reduced global segregation) in alpha band are potentially less attentive to the task at hand, and perform the task with lower accuracy. In fact, we saw that the beta band path length and alpha band connectivity across participants are negatively correlated (p < 0.05). Global segregation in both the alpha and beta bands during the encoding interval leads to higher accuracy.

### Graphical Models of Networks

Representative circos^44^ plots of the theta, alpha and beta band networks based on the appropriate median splits are shown in Figure 5. Network connections common among 4 or more participants were retained within each median split of the data. We can see that the theta band networks shows reduced clustering coefficients in the plot for slow responders (outer third circle of the figure) relative to fast responders. For alpha band networks there is a clear reduction in connections for the high accuracy participants relative to low accuracy participants. Finally the beta band network shows a reduction in long distance connections among the high accuracy participants.

## Discussion

We used graph theory to characterize the properties of networks estimated using cGGM models in different EEG frequency bands during a working memory task. During encoding increased global segregation in alpha band networks and in beta band networks predict better accuracy. Increased local integration of functional clusters in the theta band throughout the trial predicts quicker responses. Network structures present during the encoding interval likely reflect the correlates of attentional processes that affect the quality of representation in working memory^22^. Retaining a memory in the focus of attention is a process that begins in the encoding interval and continues throughout the trial as reflected by the increased local integration of functional clusters in the theta band.

### Global Segregation During Encoding Improves Accuracy

Participants who are able to reduce the influence of the background alpha band networks are able to retain memories better. This phenomenon is a function of attentional resources being directed towards the task and is likely complementary to alpha desynchronization^42^. Alpha desynchronization has been reported in tasks involving memory^23^ to positively correlate to performance, that is, greater desynchronization leads to improved performance.

In the beta band, global segregation as estimated by the path length during encoding predicts accuracy. During encoding networks become more globally separated, with greater separation particularly between frontal/temporal and all other areas. In other words, information sharing across synchronized brain areas takes longer since more (functional) distance exists between areas reducing their influence on each other. We see that greater distance particularly between frontal and parietal/occipital areas is present in the high accuracy group (see Figure 4c). This result is consistent with a recent finding that there are separate beta networks in the frontal areas and the parietal/occipital areas of the brain^45^. Increased segregation can allow for simultaneous redundant representations thereby increasing robustness of representations in the brain^7, 45, 46^. Working memory representations can be reliably decoded from both parietal and occipital areas^3^ and even from sub-regions of the frontal cortex^6^. Lesion studies suggest that patients who only have parietal lesions^46^ or patients who only have frontal lesions^45^ can in fact perform memory tasks.

### Functional Clustering and Focus of Attention

Memories in short term memory that are retained actively are believed to be in the focus of attention^47, 48^ and are available for immediate recall. During our experiment, participants needed to retain two items in memory, and in the focus of attention, throughout the trial till the retrieval interval when they needed to compare one to the probe. The changes that we see in the theta band local integration may show how this focus of attention is organized at the network level through active oscillatory regimes. This theta band network alteration is initiated during the encoding interval and sustains throughout the trial.

Changes in clustering coefficient upon the beginning of the trial differs between high and low reaction time participants (slow and fast responders). Fast responders have an increased clustering coefficient over temporal lobes during the encoding interval while slow responders do not show this change. This particular organization of changes shows that the theta band networks are involved in encoding the stimulus information as much as they are in sustaining encoded memories.

Clustering coefficient is an estimate of the local integration of functional clusters. It can be viewed as a mechanism to maintain functional homogeneity^49^ within a cluster of connections while retaining information at multiple scales^50^. For example, if areas A and B are connected to area C, information about the task that may be represented in A and B is shared also with C. We can see how the consistent functionality across the network while also maintaining information at different scales (a property of representations retained in the brain^7^) are likely important to the ability to manipulate memories rapidly.

### Simultaneous Attention and Maintenance Processes

We see that the processes facilitating maintenance of a memory (changes in local integration of functional clustering) are active from the encoding interval. Simultaneously, processes involved in attention (decreasing global segregation in alpha band and path length in beta band) are possibly involved in helping encode the stimulus information. Many studies focus on changes in network structure during the maintenance interval^18, 24, 26^ and their influence on behavior, with the interpretation that these changes are associated with processes involved in retaining a memory. Our results suggest that active network structures during the maintenance interval are manifested during the encoding interval itself.

### Graph Theory Reveals Coordination of Distributed Activity

The architecture of networks has been investigated in relation to working memory at lower scales than the EEG (i.e. neuronal level^51, 52^) showing how memories can be retained in a distributed manner through the weights of connections between nodes that can be approximated to neurons. Our work provides further evidence that the network architecture is important at multiple scales. At the scale of the EEG we see that the organization of networks in frequency bands has a robust relationship to behavior. Different network structures as revealed through graph theoretic metrics that influence performance during retrieval are present from the encoding interval itself. Future work will need to determine exactly how these higher level architectures influence lower level activity in order to retain and use active memories.

## Data Availability

The datasets generated and analysed during the current study are available from the corresponding author on reasonable request.

## Author contributions statement

R.S. conceived of the experiment, A.W. and R.S. designed the experiment, A.W. conducted the experiment and analyzed data under the supervision of R.S. All authors reviewed the manuscript.

## Additional information

The authors declare no competing interests.

